# Global prediction of candidate R-loop binding and R-loop regulatory proteins

**DOI:** 10.1101/2021.08.09.454968

**Authors:** Louis-Alexandre Fournier, Arun Kumar, Theodore Smith, Edmund Su, Michelle Moksa, Martin Hirst, Peter C. Stirling

**Affiliations:** Terry Fox Laboratory, BC Cancer, Vancouver BC, V5Z1L3; Interdisciplinary Oncology Program, University of British Columbia, Vancouver, BC; Department of Medical Genetics, University of British Columbia, Vancouver, BC; Michael Smith Laboratories, University of British Columbia, Vancouver, BC; Department of Microbiology and Immunology, University of British Columbia, Vancouver, BC

**Keywords:** R-loops, R-loop binding proteins, random forest, CRISPR, flow cytometry

## Abstract

In the past decade there has been a growing appreciation for R-loop structures as important regulators of the epigenome, telomere maintenance, DNA repair and replication. Given these numerous functions, dozens, or potentially hundreds, of proteins could serve as direct or indirect regulators of R-loop writing, reading, and erasing. In order to understand common properties shared amongst potential R-loop binding proteins (RLBPs) we mined published proteomic studies and distilled 10 features that were enriched in RLBPs compared to the rest of the proteome. We used these RLBP-specific features along with their amino acid composition to create a random forest classifier which predicts the likelihood of a protein to bind to R-loops. In parallel, we employed a whole-genome CRISPR screen coupled with flow-cytometry using the S9.6 monoclonal antibody to sort guide RNAs associated with induction of high S9.6 staining. Known R-loop regulating pathways such as splicing and DNA damage repair are highly enriched in our datasets, and we validate two new R-loop modulating proteins. Together these resources provide a reference to pursue analyses of novel R-loop regulatory proteins.

## INTRODUCTION

R-loops are nucleic acid structures consisting of an RNA:DNA hybrid in the genome and an exposed ssDNA loop on the non-template strand. R-loops are thought to primarily form co-transcriptionally although some compelling instances of R-loop formation in *trans* have been observed (1, 2). Early studies linked R-loop functions to the regulation of class-switch recombination in developing B-cells (3). However, research in the past decade has greatly expanded our understanding of the many functions of R-loops. We now know that R-loops are key regulatory intermediates of epigenetic states, telomere maintenance, DNA repair reactions, mitochondrial DNA replication, and more (4, 5). At the same time dysregulation of normal R-loop formation and dynamics is associated with genome instability, potentially impacting cancers and diseases of trinucleotide repeat-expansion (6, 7). Given the breadth of these functions and the pervasive transcription of the human genome, there is likely to be a large number of factors that regulate R-loop formation, bind R-loops, and resolve R-loops. To borrow a concept from epigenetics, the network of possible R-loop ‘writers’, ‘readers’ and ‘erasers’ is potentially very large.

Some efforts have already gone into cataloguing R-loop regulatory proteins. Physical interactions with R-loops have been assessed by immunoprecipitation with the S9.6 antibody in the presence or absence of synthetic RNA:DNA competitors followed by mass spectrometry (8). This study identified hundreds of candidate R-loop interacting proteins including many known interactors, and focused on validation of DHX9 as a novel R-loop reader in transcription termination, and eraser in genome stability maintenance (8). A related study used synthetic RNA:DNA hybrids conjugated to beads to isolate hybrid binding proteins from cell lysates (9). While both studies identified hundreds of candidate R-loop binding proteins (RLBPs) they overlapped by >100 proteins creating a higher confidence set of proteins that may preferentially recognize R-loops. Building on this set of high-confidence RLBPs it is possible to extract common protein features at the amino acid sequence level, and at a higher level such as protein domain organization or function (10–12). Based on this, here we use a random forest machine learning classifier to score R-loop binding potential for the human proteome.

Of course, binding to RNA:DNA hybrids may not correspond perfectly to factors which regulate the stability of hybrids. Proteins might bind R-loops and not affect their stability, conversely proteins may influence R-loop stability through indirect effects on transcription or DNA topology without binding R-loops. Thus, parallel functional annotation of R-loop regulators would be ideal. To date we are not aware of systematic studies to address this, however several groups have performed targeted screens for candidate R-loop regulators. A previous siRNA screen for γH2AX induction identified a large set of RNA processing factors and used RNaseH expression in a secondary screen to implicate R-loops in some of the observed DNA damage (13). More recently, work in yeast using direct S9.6 staining of chromosome spreads derived from subsets of genome instability mutants (14), or specifically DNA repair mutants (15), both found cases of genome instability linked to increased S9.6 staining. Indeed, S9.6 antibody was again used in a targeted siRNA screen against 240 DNA damage repair regulators in HeLa cells (16).

Notwithstanding specificity issues with S9.6 immunofluorescence (17), which we now know is due to cross-reactivity with ribosomal RNA (18), when used as a screening tool, S9.6 intensity enabled these and many other studies to identify new candidate R-loop regulatory proteins (15, 19–21). Here we combine a genome-wide CRISPR knockout (22) approach with S9.6 flow cytometry to identify candidate R-loop regulating genes at a large scale for the first time.

Coupling our machine learning and CRISPR screen data presents a partly-overlapping resource of candidate RLBPs and R-loop regulatory factors. While it is unlikely that all of these proteins directly influence R-loops, this dataset provides a resource for ranking candidate genes implicated in R-loop regulated processes such as telomere maintenance, RNA processing, transcription termination, DNA repair, DNA replication and the regulation of epigenetic states. Our analyses point to expanded representation for known R-loop regulatory complexes such as the spliceosome or DNA repair machinery. We also validate the association and regulatory effect on R-loops for two proteins, LIG1 and FXR1, not previously associated with R-loops. Together these analyses provide a foundation for new understanding of the varied functions of RNA:DNA hybrids in the human genome.

## MATERIALS AND METHODS

### Reference Databases

Datasets for curating the R-loop binding proteins list were taken from Cristini et al. and the R-loop attracted proteins dataset from Wang et al. (8, 9). Protein domain information was extracted from Pfam version 32.0 using an E value < 0.001 (23). Proteins longer than 5000 amino acids were excluded from this analysis. Protein abundance was mined from PaxDB 4.0 protein abundance whole-organism integrated database (24). All values from PaxDB are in parts per million. The nucleic-acid binding list was curated and compiled using DNA-binding and RNA-binding lists from Uniprot (25) and Interpro (26) supplemented with proteins identified in Beckmann et al (27) as RNA-associated to make a total of 3622 proteins. Annotated phosphorylation site data for each protein was mined from the dbPTM database (28). Protein association networks for the RLBP were created using the current STRING database version 11.0 (29). GO pathway enrichment analysis was performed using DAVID v6.8 with respective controls (30).

### Protein features and data visualization

Protein lengths and percentage of charged amino acids were calculated using in-house R scripts. The R package ‘Peptides’ was used to calculate aliphatic index and GRAVY scores were calculated by using the Kyte-Doolittle algorithm (31). Solubility scores were calculated using CamSol with default settings of pH=7.0 and proteins with non-standard amino acids were rejected (32). The overall solubility score for each sequence was used for further analysis. Disorder percentage was calculated using DISOPRED3 (33). Low-complexity region percentage was calculated using the fLPS software by predicting compositional biases for the whole protein sequence and dividing it by the total length of each protein (34). Pi-pi interaction tendency was evaluated using PScore (35). Network analysis was performed and visualized using Cytoscape (v3.8.2 (36) and GeneMANIA (v3.5.2) (37). Any additional analysis and data visualization was performed using custom R scripts (Version 3.6.3).

### Random forest classifier for R-loop binding prediction

A random forest classifier was generated using the “Caret” open-source package in R (https://cran.r-project.org/web/packages/caret/index.html) and was fed the following features for each protein - Length, Aliphatic index, GRAVY, Abundance, CamSol, Disorder and low-complexity region percentage, PScore, capability to bind to nucleic-acids, Phosphosite data, and the amino acid makeup for each of the 20 standard amino acids. Feature selection was performed using the Variable Importance evaluation function in Caret and the wrapper algorithm Boruta (38). Both analyses revealed high dependency on abundance and nucleic-acid binding capabilities of RLBPs. Using only these features, however, could result in a large number of false positives as the model would not be able to differentiate between other possible features of the protein. Feature selection was therefore not used.

For the training and testing datasets, only proteins that had values for all 30 features were chosen. 116 of the RLBP were used as the positive class while the rest of the whole proteome was chosen as the negative class. In order to avoid a class imbalance, the remaining whole proteome was randomly shuffled to choose 150 proteins at a time and 100 such shuffles were performed to avoid bias. This generated 100 different random forest models, each with its own training and testing dataset (116 RLBP + 150 different proteins). The dataset was then split into 80% training and 20% testing and the random forest algorithm was tuned using a 10-fold cross-validation approach to avoid overfitting. Additionally, the total number of features at each node (mtry) were made dependent on the highest area under the curve (ROC-AUC) to maximize ROC-AUC, thus improving prediction power. Each protein was assigned a probability score between 0 and 1 such that proteins with probabilities greater than or equal to 0.5 were classified as RLBPs. Applying an easy ensemble approach, the probabilities from all 100 models were averaged for each protein and subjected to thresholds. Proteins with an average probability > 0.7 over 100 models were extracted for higher confidence resulting in a final set of 669 proteins. Scripts used for RF tuning, development and prediction can be found on Github (https://github.com/arunk95/RLBP_Prediction).

### Cell Culture, siRNA transfection and drug treatments

HEK293T/17 were cultivated in Roswell Park Memorial Institute Medium (RPMI 1640) supplemented with 10% FBS, 1% Pen/Strep, 10µM HEPES pH 8.0, 50µM 2-β-mercaptoethanol, 1mM sodium pyruvate and 2mM GlutaMAX. HeLa cells were cultivated in Dulbecco’s modified Eagle’s medium (DMEM) (Stemcell technologies) supplemented with 10% fetal bovine serum (Life Technologies) in 5% CO2 at 37°C. For RNA interference, cells were transfected with siGENOME-SMARTpool siRNAs from Dharmacon (Non-targeting siRNA Pool #1 as si-Cont, ON-TARGETplus Human LIG1 3978 siRNA siRNA SMARTpool and ON-TARGETplus Human FXR1 8087 siRNA SMARTpool). Transfections were performed with Dharmafect1 transfection reagent (Dharmacon) according to manufacturer’s protocol and harvested 48 h after siRNA transfection. For drug treatments, cells were treated with 5µM tigecycline (Millipore Sigma, PZ0021) for 72 hours prior to fixation, or 25µM L82-G17 (Aobius, AOB31453) for 4 hours prior to fixation.

### TKOv3 library cloning and viral preparation

The Toronto human knockout pooled library (TKOv3) was a gift from Jason Moffat (Addgene #90294). The library was amplified in bacteria as described in the Moffat protocol on Addgene (https://www.addgene.org/pooled-library/moffat-crispr-knockout-tkov3/). The amplified sgRNA library was packaged into lentiviral particles using HEK293T/17 cells by co-transfection with psPAX2 and pMD2.G viral plasmids (1:1:1 molar ratio). To produce viral particles at scale, 510×10^6^ cells were transfected in 60 15-cm culture dishes. Each dish was transfected using the TransIT-LT1 Transfection reagent (Mirus, MIR 2305, 3µL/µg of DNA) with 4.8μg psPAX2, 3.8μg pMD2.G and 8μg library DNA 24-hours after cells were seeded. After 24 hours of transfection, the media was replaced with high-BSA growth media (DMEM + 1.1g/100mL BSA + 1X Pen/Strep) for viral harvest. Lentiviral particles were harvested twice at 48-hours and 72-hours post-transfection. Both harvests were pooled, viral particles were concentrated by ultracentrifugation (25,000g) and frozen at −80°C for future use.

Viral titre was assessed to determine the multiplicity of infection (MOI) of the concentrated viral particles. HeLa cells were seeded at a density of 2.5×10^6^ cells/well in 12-well plates and spin-fected (2000 rpm for 2 hours at 37°C) with increasing concentrations of virus in the presence of 8µg/mL polybrene (Millipore TR-003-G lot#3287963). The next day, the cells were split into 6-well plates at a density of 0.5×10^6^ cells/well and subjected to 2µg/mL puromycin (Sigma P8833) for selection of infected cells. Cells not transduced with the library (no virus control) did not survive past 24 hours of puromycin selection (2µg/mL). After 48-hours of selection, all cells in all wells were counted to determine the viral volume that resulted in 20-40% survival in puromycin (corresponding to an MOI of 0.2-0.4 assuming an independent infection rate).

### CRISPR knockout pool generation

The FACS-based CRISPR screen was performed in biological duplicates. HeLa cells were transduced with sgRNA libraries at a multiplicity of infection of (MOI) of 0.3, aiming for coverage of, on average, 600 cells per sgRNA reagent (600X). For the genome-wide screen, 300 million HeLa cells were transduced in 12-well plates (2.5×10^6^ cells per well) using the appropriate volume of viral particles for MOI of 0.3 and 8µg/mL polybrene. Twenty-four hours after transduction, the cells were trypsinized and plated into 15-cm dishes at a density of 8×10^6^ cells per dish in media containing 2µg/mL puromycin. Control 6-well plates were also plated at 0.5×10^6^ cells/well to confirm appropriate multiplicity of infection (MOI).

Forty-eight hours post-selection with puromycin, the cells were split and MOI was assessed. Cells were maintained in culture and split as needed to ensure confluence did not exceed 90%. Three days post-selection (D3), the cells were harvested for subsequent analysis by flow cytometry.

### Flow cytometry

The flow cytometry staining protocol was adapted from Forment and Jackson(39). Cells harvested for flow cytometry (∼380×10^6^ cells) were washed in PBS and resuspended at 15×10^6^ cells/tube in 15-mL Falcon tubes. Cells were spun* for 3 minutes at 4°C, the supernatant was removed and cells were resuspended in 15mL 1X PBS. Cells were centrifuged again, supernatant was removed and cells were resuspended in 1.5mL CSK buffer (25mM HEPES ph7.4, 50mM NaCl, 1mM EDTA, 3mM MgCl_2_, 300mM sucrose, 0.5% vol/vol Triton X-100 and complete protease inhibitor cocktail tablet) for 15 minutes on ice. Cells were then washed with 14mL PBS-B (1mg/mL BSA solution in 1X PBS) and spun. The supernatant was removed and cells were fixed using 1.5mL Fixation Buffer (2% wt/vol paraformaldehyde in 1X PBS) for 30 minutes at RT while rotating. The cells were subsequently washed with 8mL PBS-B, spun and stored overnight at 4°C in 1mL Storage Buffer (3% vol/vol heat-inactivated FBS and 0.09% wt/vol sodium azide solution in 1X DPBS).

The samples were washed with 8mL 1X PBS, spun and stained with primary antibody (Kerafast anti DNA-RNA hybrid [S9.6] Ab, cat: ENH 001, Lot: 071718_3, 1:200) in 1X BD FACS Wash Buffer (BD Biosciences) overnight at 4°C while rotating. Note that a sample of unstained cells was collected prior to primary antibody staining to use as a control for the flow cytometry gating. These cells were stained with secondary antibodies later on (i.e. no primary antibody control). Following the primary antibody staining, the cells were washed with 5mL 1X Wash Buffer, spun and incubated with secondary antibody (AlexaFluor Mouse 568, 1:1000) for 30 minutes at room temperature (RT) in the dark. The cells were then washed with 1X Wash buffer and resuspended at 7-10×10^6^ cells/mL in Analysis Buffer (0.02% wt/vol sodium azide in PBS-B). The cells were transferred into filtered-cap flow tubes and incubated at 37°C for 30 min before sorting. For each replicate, about 42 million cells total (600X coverage) were sorted into three populations based on S9.6 intensity: Bottom 10%, Middle 80% and Top 10% (**Figures 4A** and **S4B**).

^*^All centrifugation steps were carried out at 400g for 3minutes at 4°C.

### Genomic DNA extraction

Genomic DNA extraction was performed according to the method described in Chen and Sanjana et al. (40).

### Library preparation and sequencing

The sgRNA libraries were prepared using two-steps of PCR as described in the Moffat protocol (https://media.addgene.org/cms/filer_public/44/9f/449f22c6-f818-4af6-8af4-2bab1d20f00a/moffat_lab_crispr_screen_protocol_v2.pdf): 1) amplify the sgRNA region within genomic DNA; 2) amplify sgRNAs with Illumina TruSeq adapters with i5 and i7 indices (see **Table S5**). These indices are unique sequences that are added to DNA samples during library preparation and act as sample identifiers during multiplex sequencing. All PCR steps were performed using the NEBNext Ultra II Q5 Master Mix high-fidelity polymerase. For PCR #1, the thermocycling parameters were 98°C for 30s, 25 cycles of (98°C for 10s, 66°C for 30s, 72°C for 15s), and 72°C for 2 minutes. In each PCR #1 reaction, we used 3.5μg of gDNA. For each sample, the necessary number of PCR #1 reactions was used to capture the appropriate representation of the screen. For instance, assuming a diploid genome is ∼7.2 pg and one guide-RNA per genome, 100 µg of genomic DNA yields ∼200-fold coverage of the TKOv3 library. In this example, at least 29 PCR#1 reactions were necessary to capture 200X representation of the library for each replicate.

PCR #1 products were pooled for each biological samples and 5 µL was used for amplification and unique barcoding in PCR #2. The thermocycling parameters for PCR#2 were 98°C for 30s, followed by 10 cycles of (98°C for 10s, 55°C for 30s and 65°C for 15s), and 65°C for 5 minutes. The PCR #2 products were validated using gel electrophoresis and purified using the QIAquick Gel extraction kit (QIAGEN). The purified libraries were then quantified by Nanodrop and Qubit prior to sequencing on an Illumina MiSeq.

### Amplicon scoring and guide RNA enrichment

Amplicon sequencing data was processed for sgRNA representation using custom BASH scripts. To summarize, the sequencing reads were first de-multiplexed using the 8-bp barcodes in the forward primer, then using the 8-bp barcodes in the reverse primers. De-multiplexed reads were trimmed to leave only the 20-bp spacer (sgRNA) sequences. The sgRNA sequences were then mapped to the reference TKOv3 library sgRNA sequences using Bowtie 2(41). For mapping, no mismatches were allowed, and mapped sgRNA sequences were quantified by counting the total number of reads.

Differential fold-change estimates between samples were generated for each sgRNAs using DESEQ2 (42) by comparing enrichment between the S9.6 “high” and “low” fractions (top 10% vs bottom 10%). For the purpose of assigning probability values to the CRISPR hits presented in the network shown in Figure 5, each guide was assigned a Z-score calculated as follows (see **Table S4)**:

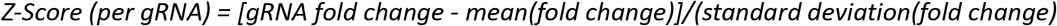

### Proximity ligation assay

Proximity ligation assay (PLA) experiments were performed using the Duolink PLA kit (Millipore Sigma). For all PLA experiments, cells were grown on coverslips before fixing. For S9.6 staining, cells were fixed with ice-cold methanol for 10 minutes and permeabilized with ice-cold acetone for 1 minute. After permeabilization, cells were washed with PBS and blocked for 1 hour at RT in Duolink Blocking Solution. Cells were then incubated with primary antibodies (S9.6 Millipore MABE1095 + anti-LIG1 Abcam ab177946 or anti-FXR1 Abcam ab155124) diluted in Duolink Antibody Diluent overnight at 4°C. Cells were washed twice (5 min) in µL/ cover slip, 1:4 PLA probe in PLA Antibody Diluent). Cells were washed (2x 5 min) with PLA Wash Buffer A, after which they were incubated 30 minutes at 37°C with PLA ligation mix (15µL/cover slip, 1:40 PLA ligase 40X in 1:5 ligation buffer 5X in Ultra H2O). Cells were washed (2x 2 min) with PLA Wash Buffer A and incubated 100 minutes at 37°C with PLA Amplification mix (15µL/slide, 1:80 polymerase solution in 1:5 amplification stock in ultra H2O). The cells were washed (2x 10 min) in PLA Wash Buffer B. After an additional wash (1 minute) in PLA Wash Buffer B 0.01X, the slides were mounted in Duolink Mounting Media with DAPI. Imaging was performed on a Leica Dmi8 microscope at 100X. ImageJ was used for image processing and quantification (43). For *in vitro* RNaseH, RNaseIII and RNaseT1 treatment, cells were treated with RNaseH (New England Biolabs, M0297S) for 2 hours, or ShortCut RNaseIII (New England Biolabs, M0245S) or RNaseT1 (ThermoFisher Scientific, EN0541) for 30 minutes at 37°C after permeabilization before blocking.

### Immunofluorescence

For all immunofluorescence experiments, cells were grown on coverslips overnight before transfection with 50µM siRNA (siCTRL, siLIG1 or siFXR1). Forty-eight hours post-transfection, the cells were fixed with ice-cold methanol for 10 minutes and permeabilized with ice-cold acetone for 1 minute. After permeabilization, cells were washed with PBS and blocked in 3%BSA, 0.1% Tween 20 in 4X saline sodium citrate buffer (SSC) for 1 hour at RT. Cells were then incubated with primary antibody (S9.6, Millipore) overnight at 4°C. Following 2 PBS washes and an additional 20 minute blocking step, the cells were then incubated with Alexa-Fluor-568-conjugated secondary antibodies for 1 hour at RT. After washing 3 times with PBS, the cells were stained with DAPI before mounting and imaging on a LeicaDMI8 microscope at 100X. ImageJ was used for image processing and quantification of nuclear S9.6 signal (43). For *in vitro* RNaseH, RNaseIII and RNaseT1 treatment, cells were treated according to the manufacturer’s guidelines with *in vitro* RNaseH (New England Biolabs, M0297S) for 2 hours, or ShortCut RNaseIII (New England Biolabs, M0245S) or RNaseT1 (ThermoFisher Scientific, EN0541) for 30 minutes at 37°C after permeabilization before blocking.

### Western blotting

Whole-cell lysates were prepared with RIPA buffer containing protease inhibitor (Sigma) and phosphatase inhibitor (Roche Applied Science) cocktail tablets and the protein concentration were determined by Bio-Rad Protein assay (Bio-Rad). Equivalent amounts of protein were resolved by SDS-PAGE and transferred to polyvinylidene fluoride microporous membrane (Millipore), blocked with 1.5% BSA in H20 containing 0.1% Tween-20 (TBS-T), and membranes were probed with the following primary antibodies: anti-LIG1 (Abcam ab177946) (1:1000), anti-FXR1 (Abcam ab155124) (1:1000) and anti-GAPDH (Thermo Scientific MA5-15738) (1:3000). Secondary antibodies were conjugated to horseradish peroxidase (HRP) and peroxidase activity was visualized using Chemiluminescent HRP substrate (Thermo Scientific).

### DRIP and ChIP qPCR

DRIP was performed using the SimpleCHIP enzymatic chromatin IP kit (Cell Signaling Technologies) in accordance with manufacturer’s instructions with some modifications. HeLa cells were seeded at 1×10^6^ in 10 cm plates and transfected the following day with siRNAs (siCTRL, siLIG1, siFXR1). Forty-eight hours after transfection, the cells were crosslinked in 1% formaldehyde for 10 minutes, followed by 2 minute incubation in 1 x glycine solution (CST#7005). Cells were washed in 10 mL ice-cold 1X PBS twice and scraped into 0.5 mL ice-cold 1X Buffer A (with 500 µM DTT and 1 x protease inhibitor cocktail (PIC) (CST#7012) per IP prep in sonication tubes. Nuclei were pelleted and resuspended in 150 µL of 1 x ChIP buffer and 1 x PIC, followed by incubation on ice for 10 minutes. DNA was sonicated on Q Sonica Sonicator Q700 for 8 min (30 sec ON, 20 s OFF) at 100 µm amplitude to fragment DNA. Lysates were clarified with centrifugation and the supernatant was collected DRIP and stored at −80°C until further use. DNA samples for DRIP were treated with RNAse A (CST#7013) followed by proteinase K (CST#10012) overnight at 65°C. DNA samples were purified with Cell Signaling spin columns (#14209S). DNA concentration was measured using NanoDrop One/Onec. DNA samples were used for DRIP analysis, where 2% input sample was isolated and DNA was separated into untreated and RNaseH-treated groups. 1 μL of 5000 units/mL RNaseH (NEB, M0297L) was used per µg of DNA and incubated for 48 hours at 37°C. ChIP-Grade protein G magnetic beads (Cell Signaling, 9006S - 25 μL per IP) were pre-blocked in pre-blocking buffer (PBS/EDTA containing 0.5% BSA) for 1 hour rotating at 4°C. Beads were immobilized with S9.6 antibody (MABE1095) using 5 µg per IP, in 1 x ChIP buffer and 1 x PIC for 4 hours rotating at 4°C. DNA was then added to the beads/Ab complex and incubated overnight with rotation at 4°C. Samples were washed 3 times with 1mL low salt buffer (0.1% SDS, 1% TritonX-100, 2mM EDTA pH8.0, 20mM Tris-Cl pH 8.0 and 150mM NaCl), 1 time with 1mL high salt buffer (0.1% SDS, % TritonX-100, 2mM EDTA pH 8.0, 20mM Tris-Cl pH 8.0 and 500mM NaCl), and eluted in 80µL of 1x elution buffer (CST#7009) for 30 minutes at 65°C vortexing intermittently. Beads were pelleted and supernatant was purified with Cell Signaling spin columns (CST#14209S). DRIP-qPCR primer sequences can be found in **Table S5**. qPCR was performed on AB Step One Plus real-time PCR machine (Applied Biosystems) using Fast SYBR Green Master Mix (Applied Biosystems).

## RESULTS

### Defining characteristics of consensus R-loop interacting proteins

We set out to understand features of R-loop binding proteins by analyzing their protein sequences and higher-order features. To develop a consensus dataset, we intersected two studies that document R-loop interacting proteins using orthogonal enrichment approaches and mass spectrometry. Upon intersecting the Cristini et al. dataset (469 proteins) and Wang et al. dataset (313; subset of proteins attracted to R-loops) we find a high confidence dataset of 122 proteins that preferentially bind R-loops that we classify as R-loop binding proteins (RLBPs) (**Fig. 1A**)(**Table S1**) (8, 9). We performed a Pfam domain enrichment analysis on the RLBPs and as expected they were enriched for multiple RNA-binding domains including the RRM motif, OB-fold domains and the DEAD/DEAH-box domains that have previously been linked to R-loop binding proteins (**Fig. 1B**) (8, 9). Additionally, RLBPs are significantly enriched for proteins with three or more domains compared to the whole human proteome and the nuclear proteome (**Fig. 1C**). While all datasets share similar proportions of proteins with two domains, the RLBP dataset has relatively few proteins containing one or no domain indicating that RLBPs have a high valency, in that RLBPs can probably form multiple protein-protein interactions. This is further supported by a tightly interconnected protein-protein physical interaction network of RLBPs reported by the STRING database (p value < 1.0e^-16^; https://string-db.org/). Since RLBPs must recognize the RNA or DNA region of the R-loop, we hypothesized the RLBPs should be enriched for nucleic-acid binding domains. Indeed, when compared to both the whole proteome and even the nuclear proteome there was a strong enrichment for nucleic-acid binding domains, supporting that this is an important property of RLBPs (**Fig. 1D**).

**Figure 1.**
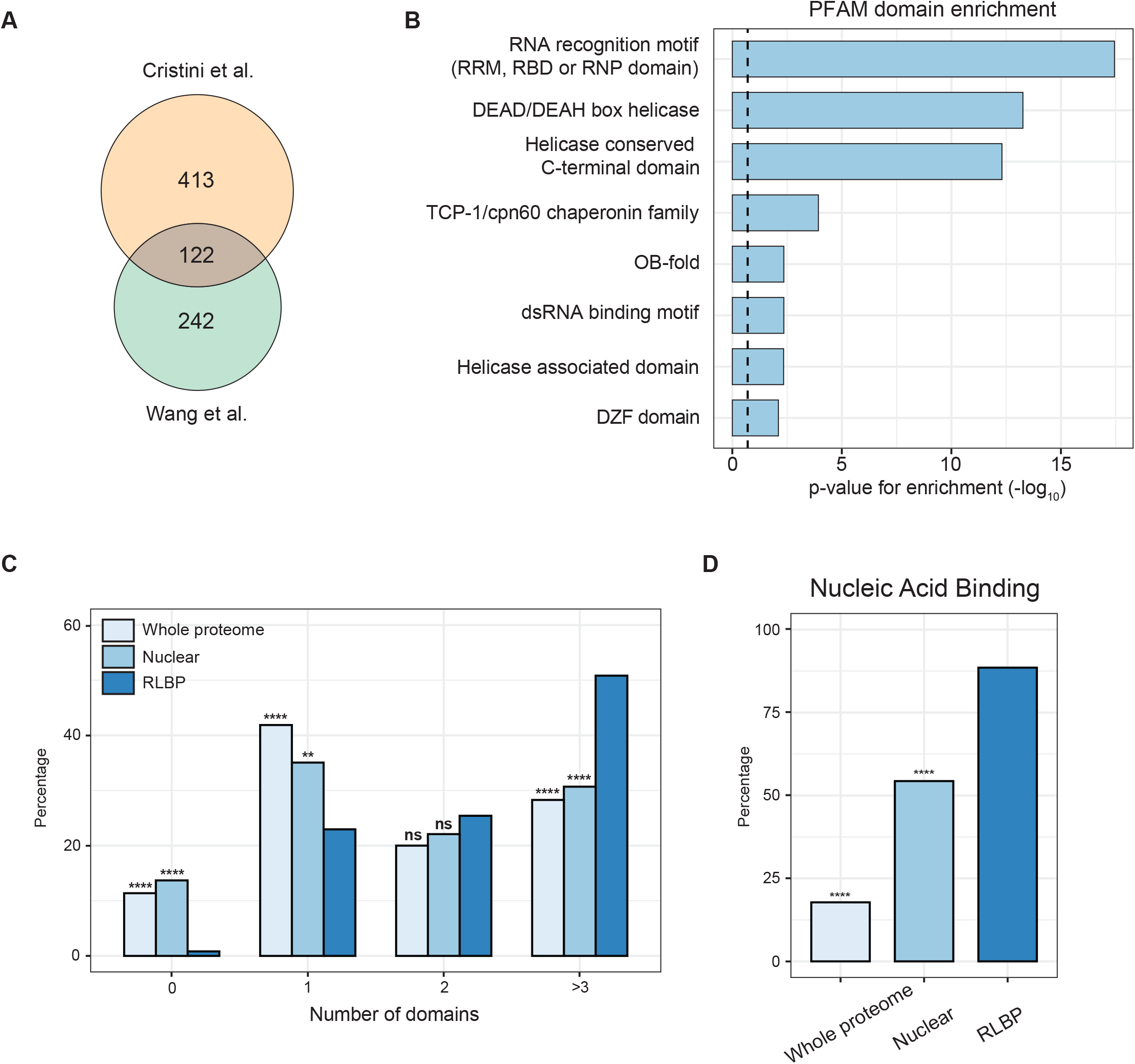
Higher order features of R-loop binding proteins. (A) Venn diagram of candidate R-loop binding proteins identified in published studies. (B) Enrichment analysis of PFAM domains enriched within the intersected RLBPs (False discovery rate adjusted p-value < 0.01) (C) Comparison of domain numbers between the whole proteome, nuclear proteome and RLBP datasets. (D) The proportion of proteins in each category encoding a nucleic acid binding domain. For C and D, ****p<0.0001, **p<0.01, ns p>0.1 Fisher’s exact test.

### Protein sequence and functional characteristics of the R-loop binding proteome

Next, we investigated the potential for common protein sequence features of the RLBPs. First assessing protein length we found that the RLBPs are significantly longer than the average human protein (**Fig. 2A**). We then assessed the hydrophobic character of each protein by calculating GRAVY (Grand Average of hydropathy) scores (31). On average, the RLBPs have lower GRAVY scores versus the human proteome suggesting a more hydrophilic character that could aid in a more flexible protein structure and associate with nucleic acid binding properties (**Fig. 2B**). Interestingly, the difference between the RLBP and the nuclear proteins was not significant suggesting that this property might be common to all nuclear proteins and not just the RLBPs. Consistent with low GRAVY scores, when we calculated the aliphatic index we found that the RLBPs have significantly lower indices compared to the rest of the proteome (**Fig. S1A**). Surprisingly, RLBPs have a higher average aliphatic index than a nuclear control indicating that the RLBPs might have a characteristic range of value that could be used to differentiate them from other nuclear proteins. We next investigated the amino acid makeup of the RLBPs and found that they are enriched for charged amino acids (**Fig. 2C**). Interestingly, this trend holds true for both positive and negatively charged amino acids (**Fig. S1B-C**). Overall, we find that a typical RLBP is longer, more stable, more hydrophilic and enriched for charged amino acids when compared to the rest of the human proteome which might prime them for interactions with other proteins.

**Figure 2.**
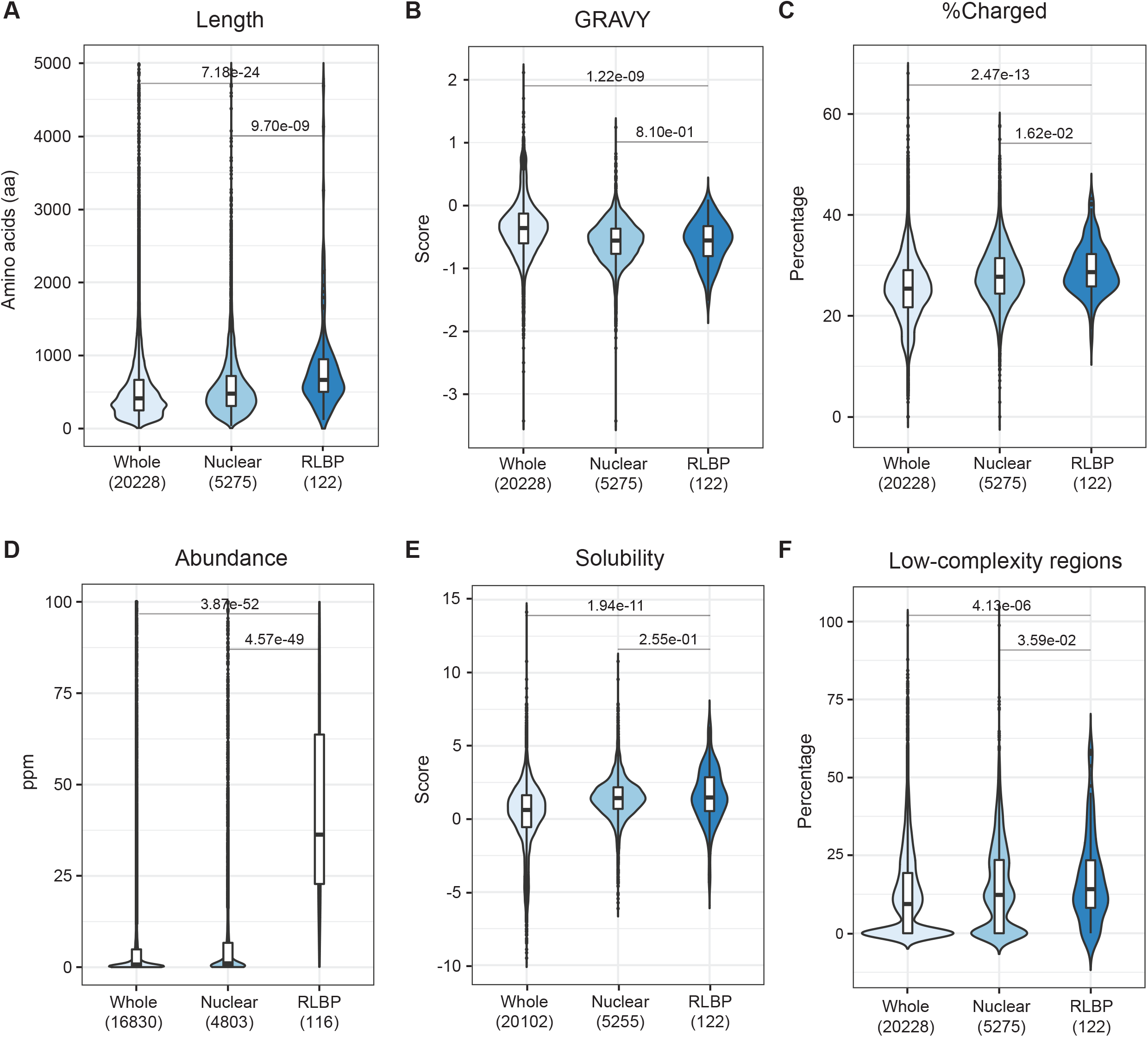
Amino acid sequence characteristics of R-loop binding proteins. (A-F), Length (A), hydrophobicity and hydrophilicity (GRAVY) (B), percentage of charged residues (C), protein abundance(D), solubility (E), and percentage of low-complexity regions (F) were compared for the indicated whole proteome, nuclear proteome and RLBP datasets. The exact p-values resulting from a Mann-Whitney-Wilcoxon test after Bonferroni correction for multiple comparisons are reported.

We next sought to characterize the biochemical properties of the RLBP dataset. We first compared protein abundance using the PaxDB database (24) and found that RLBPs are more abundant than the average of both whole proteome and nuclear proteins (**Fig. 2D**). We then measured solubility scores using the CamSol solubility prediction method that utilizes protein sequences to assign solubility scores (32). We find that the RLBPs have significantly higher scores indicating they are more soluble and may have a lower tendency to form aggregates (**Fig. 2E**). Interestingly, RLBPs have similar CamSol scores to all nuclear proteins indicating that this is not a distinct feature of the RLBPs but a feature of nuclear proteins. So far RLBPs appear to be abundant, soluble, charged proteins, and we wondered if this would affect their propensity to phase-separate. Intrinsically disordered regions (IDRs) and low-complexity regions (LCRs) in proteins have been implicated as key drivers of liquid-liquid phase separation (44).

These regions do not form canonical three-dimensional structures making them more flexible and prone to interactions. Previous studies have noted the presence of IDRs in some RLBPs (9, 45), and accordingly DISOPRED3 disorder scores show that RLBPs are more disordered than the human proteome but not significantly different from nuclear proteins (**Fig. S1D)** (33). The same trend held true while measuring propensity to phase separate through ***π***-***π*** interactions **(Fig. S1E**) (35). We also measured the presence of LCRs using fLPS which annotates compositional biases in a given sequence to provide a total number of LCRs in a protein (34). As with protein disorder, RLBPs present a significant enrichment of LCRs compared to the human proteome (**Fig. 2F**). Finally, we used phosphorylation as a proxy to determine the proportion of the proteome that could be post translationally modified (**Fig. S1F**) (28). Overall, we find that the RLBP dataset is more abundant, soluble, disordered, has a higher valency and greater tendency to be post translationally modified compared to the rest of the proteome.

### A random forest classifier of R-loop binding properties in the human proteome

Our data so far suggests that high confidence RLBPs have common sequence features, functional properties and domain architectures. Thus, we hypothesized that these features could be used to predict new candidate RLBPs in the rest of the human proteome. Using the features of RLBPs we set out to develop a tool that can assign a score of the likelihood that any given protein in the proteome is an R-loop binding protein. We employed a machine learning approach using the Random Forest (RF) algorithm to develop a binary classifier that could yield predictions of the highest confidence. The RF algorithm is a common machine learning method owing to its versatility, and its ability to leverage big data with multiple variables (46). In total, our RF model assesses 30 features including: Length, GRAVY, aliphatic index, abundance, CamSol scores, disorder%, LCR%, PScore, ability to bind nucleic-acids, Phosphosite%, and the amino acid makeup of all 20 standard amino acids (**Table S4**). We chose not to eliminate any features after performing feature selection to reduce possible false positives (**see Methods**).

To train and test our classifier, we chose the 122 overlapping RLBPs as our positive class while the rest of the proteome made up the negative class. To avoid class imbalance, we employed an easy ensemble approach where we randomly shuffled the negative class 100 times and chose 150 proteins from each iteration, resulting in 100 different training and testing datasets and therefore 100 different RF models. Using an 80% training, 20% testing split, along with 10-fold cross-validation, we assessed the performance of our classifiers (**Fig. 3A**). On average, our classifiers achieved an accuracy of 91.6%, an F1 score of 0.893 and a Matthews correlation coefficient (MCC) of 0.827 on the test set (**Fig. 3B** and **Table S3**). Each protein was assigned a probability score between 0 and 1, where proteins greater than or equal to 0.5 are classified as candidate RLBPs. To determine a probability cut-off for new candidate RLBPs, we plotted the density function for both the RLBPs and the remaining proteome (**Fig. 3C**). The RLBPs peak at 0.95 whereas the remaining proteome peaks at 0.05. This was not surprising as the RF model should predict RLBPs as positive (i.e. ∼1) and the rest of the proteome as negative (i.e. ∼0).

**Figure 3.**
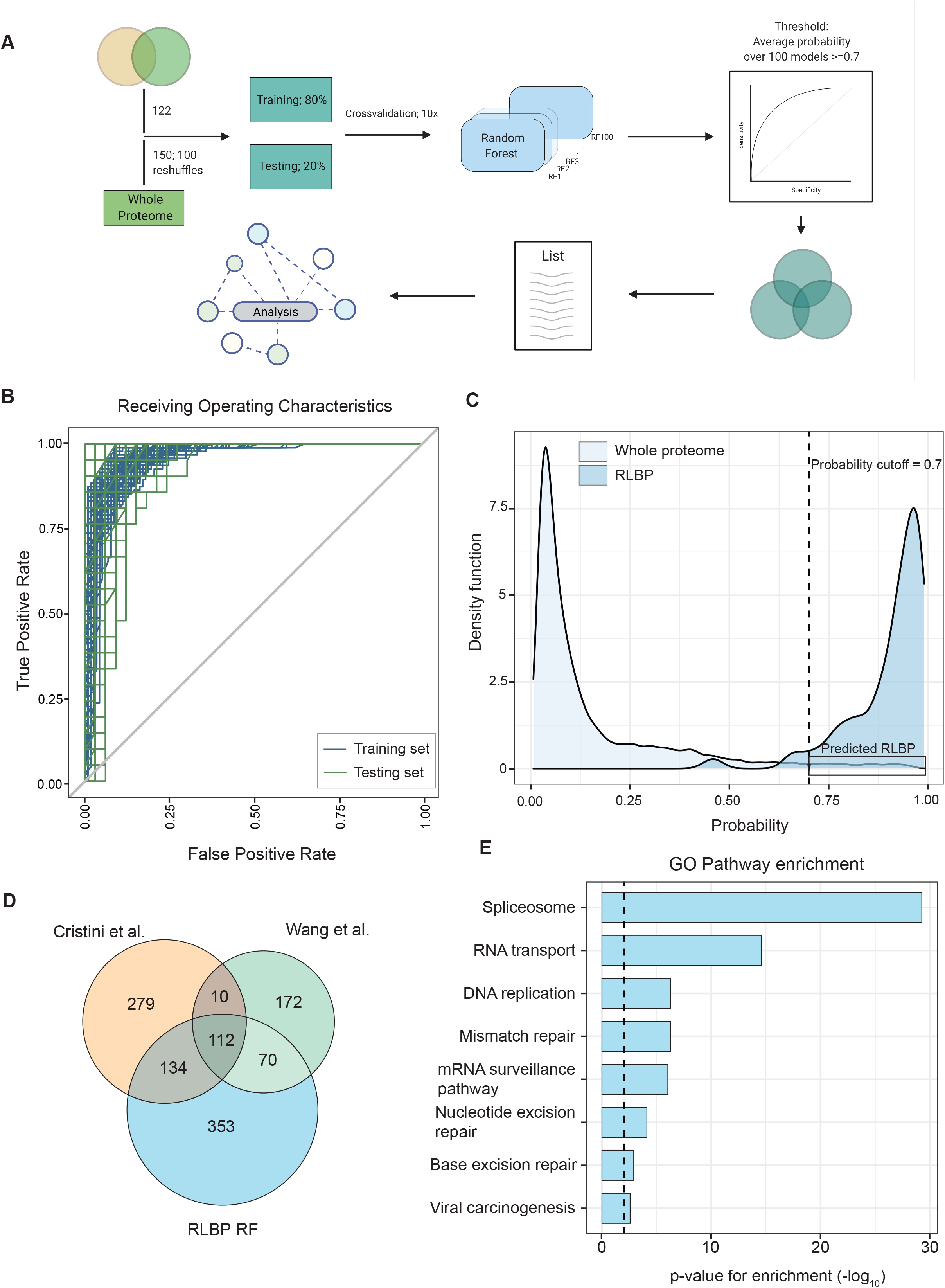
A random forest classifier to determine R-loop binding property likelihood. (A) Schematic of the machine learning model pipeline (created with BioRender). (B) Receiving operating characteristics of the training (blue) and testing (green) sets. (C) Whole proteome prediction of RLBP character by the model. The plot shows the probability distribution of the whole proteome and our RLBP training set. The newly predicted RLBPs extracted from the whole proteome are highlighted on the right. (D) Venn diagram of overlap between published datasets and the random forest predicted RLBPs at a probability cut-off of 0.7. (E) Enriched pathways upon gene ontology analysis for the 557 uniquely predicted proteins (False discovery rate adjusted p-value < 0.01).

Importantly, the tail end of the whole proteome set falls in a higher probability range indicating newly predicted RLBPs (rectangular box in **Fig. 3C**). Since both distributions intersect at 0.65, we chose 0.7 as a conservative threshold to call new candidate RLBPs. Using 0.7 as our cut-off, we present a set of 669 proteins that are predicted as R-loop binding by the RF algorithm (**Fig. 3D**). When intersected with the published datasets used to produce the original list of 122 proteins, the averaged RF models predict 112 of the 122 proteins used to train the model. Of the remaining 557 proteins, the model successfully predicts 134 proteins found in the Cristini dataset and 70 found in the Wang dataset (8, 9). This represents a significant enrichment (p=6.0e^-117^ Cristini; p=1.4e^-59^ Wang, Fisher’s exact test) suggesting there are indeed common properties of RLBPs that can be extracted from the proteomic datasets. These 206 proteins represent a second tier of candidate RLBPs that are both predicted by our feature model, and found in one published proteomic study (**Fig. 3D**). Additionally, our model predicts 346 proteins not previously identified as a resource of novel candidate RLBPs. Gene Ontology analysis for pathway enrichment on the same set of proteins revealed strong enrichment for proteins involved in spliceosome formation and function, RNA transport, DNA replication and various forms of DNA repair (**Fig. 3E**). Most of these processes have been implicated in R-loop biology by previous studies (4, 5, 47). To determine whether nucleic acid binding overly skews the RF algorithm, we compared the assessed features between the predicted RLBPs and the remaining nucleic acid binding proteins. Except for GRAVY and LCRs, there is a significant enrichment for all remaining features supporting our hypothesis that the RLBP dataset is a unique subset of cellular proteins with potential R-loop binding ability (**Fig. S2A-I**). For independent validation, we queried the scores of proteins found in an independent S9.6 based mass spectrometry study (48). Our model predicts 20 of the 33 proteins unique to this independent validation dataset (p=4.2e^-27^ Fisher’s exact test) (**Fig. S3A**). Together our model recovers the vast majority of high confidence RLBPs, selects new candidates found in previous mass spectrometry studies, and assigns an RLBP probability to the rest of the human proteome.

### Genome-wide CRISPR-flow cytometry screening for regulators of S9.6 antibody staining

We predicted that a subset of the RLBPs will have an impact on R-loop stability and function. While we cannot hope to individually validate each of the new RLBPs here, an ideal scenario would be to use high-throughput approaches to look for genes whose disruption alters R-loop levels in cells. Previous studies on a small scale have identified proteins that can affect R-loop stability using the S9.6 monoclonal antibody (14–16). Other studies have used DNA damage and RNaseH1 expression as a proxy for R-loop accumulation and associated genome instability (13). Antibodies that recognize nucleic acids in a non-sequence specific way are challenging as they can cross-react with other structures. While S9.6 clearly recognizes DNA:RNA hybrid structures in R-loops, these hybrids of course can appear elsewhere (e.g. okazaki fragments, resected DNA ends, retrotransposons, etc.), and S9.6 cross reacts with structured RNAs, especially ribosomal RNA (18). Thus, while increases in S9.6 staining can indicate changes in other cellular RNAs more broadly, the staining also can indicate changes in DNA:RNA hybrid levels associated with R-loops. As a first step to implementing a high-throughput analysis of S9.6 staining, we tested whether known R-loop inducing stresses can increase S9.6 staining by flow cytometry. Accordingly we treated HeLa cells with the topoisomerase inhibitor camptothecin and this led to a clear quantifiableincrease in S9.6 staining by FACS (**Fig. S4A**).

Since we can use FACS to differentiate high S9.6 staining from low S9.6 staining, we applied this approach to a large pool of HeLa cells bearing the TKOv3 genome-wide CRISPR guide RNA library (22). We infected 3e^8^ HeLa cells with the TKOv3 library at a multiplicity of infection (MOI) of 0.3 to attain 600X coverage, selected the infected cells with puromycin over 48 hours prior and outgrew the population over three days. The cells were then stained with the S9.6 antibody for flow cytometry analysis and sorted into three populations based on S9.6 intensity: the top 10%, a bulk fraction (middle 80%) and the bottom 10% (**Fig. 4A** and **S4B**). Deep amplicon sequencing of the guide regions extracted from genomic DNA purified from each population enabled us to evaluate sgRNA enrichment between groups using a differential gene expression approach comparing the top and bottom populations using DESEQ2 (42, 49). This analysis uncovered a consensus list of 445 enriched sgRNAs (420 unique sgRNAs) in the R-loop “high” population, representing potential R-loop modulating factors (**Fig. 4B**). Since the TKOv3 library contains 4 sgRNAs for every gene it targets, we were surprised that the differential analysis did not significantly enrich multiple sgRNAs per gene. This can likely be explained by the stringent gating strategy utilized (i.e. top 10%) to limit the occurrence of false positives, which allowed us to identify genetic disruptions with strong phenotypic consequences on S9.6 staining, perhaps at the expense of milder phenotypes. Therefore, sgRNAs whose distributions did not fall within the top 10% of S9.6 staining did not come up as enriched in our analysis. Nevertheless, this approach identified known R-loop modulators along with novel candidates (see below). Gene ontology analysis of the biological processes represented within this list revealed a surprising enrichment of mitochondrial proteins, as well as known R-loop regulatory pathways such as telomere maintenance or DNA damage response (**Fig. 4C**).

**Figure 4.**
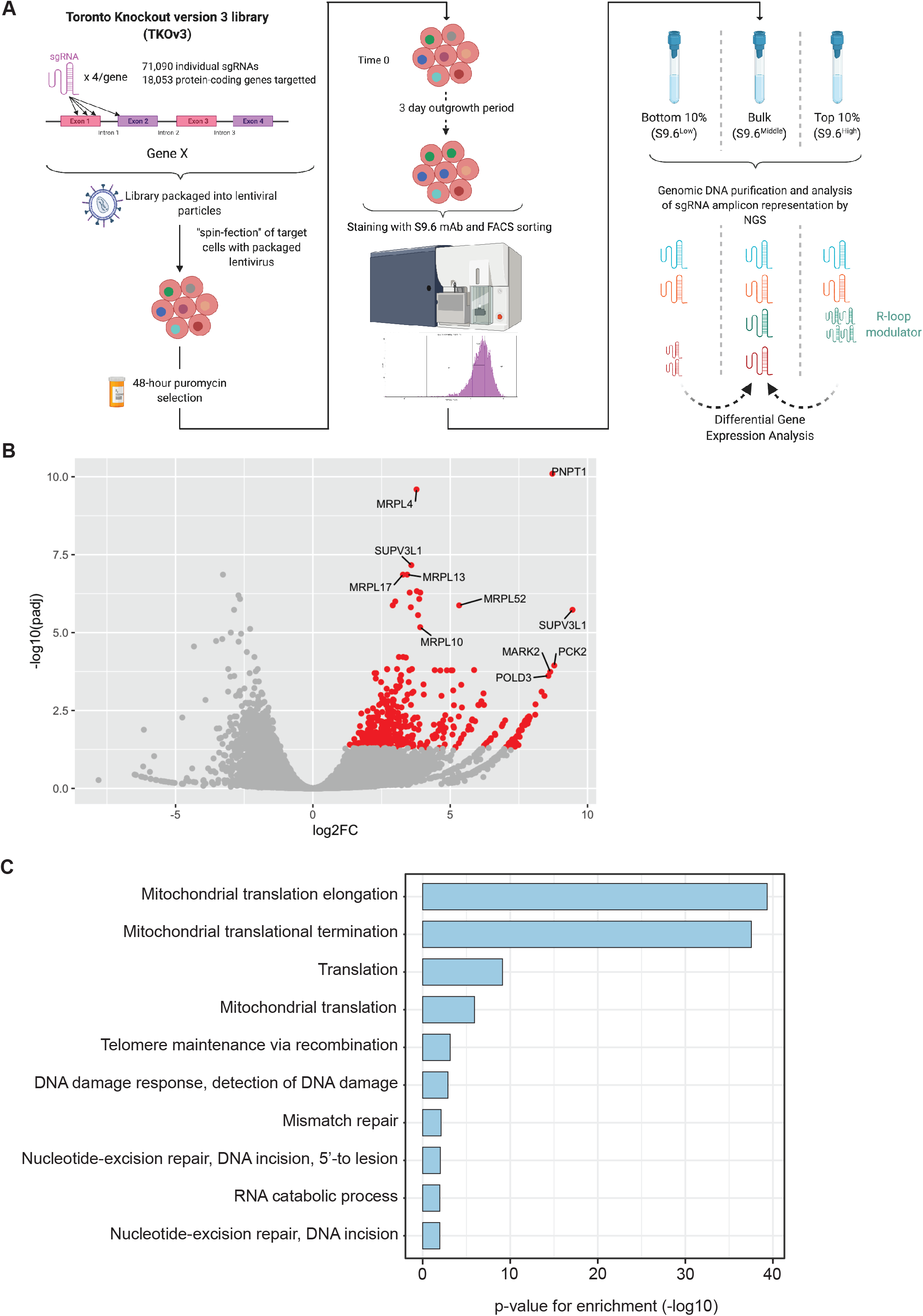
Genome-wide CRISPR screening identifies modulators of S9.6 antibody staining. (A) Schematic of CRISPR screening workflow (created with BioRender). (B) Plot of log2FC vs -log10(padj) of differentially expressed sgRNAs in the S9.6 “high” vs S9.6 “low”population. Significantly enriched sgRNAs in the S9.6 “high” group are highlighted in red (padj < 0.05). The top 10 most significantly enriched sgRNAs are labelled. (C) Top 10 enriched biological processes represented in the gene ontology analysis of FACS screen hit list (David v6.8, -log10[p-value]).

### The functional spectrum of candidate RLBPs

With the machine learning and CRISPR-FACS screens in hand we undertook a network analysis to identify cellular complexes enriched by the datasets (**Fig. 5**). Protein complex analysis on the newly predicted 557 proteins by our RF classifier combined with the genome-wide CRISPR screen gene list revealed the presence of 10 significantly enriched complexes (p<0.01)(**Fig. 5A**). These include the C complex spliceosome (50), the 55S mitochondrial ribosome (51), the genome surveillance complexes RAP1, BASC, and 53BP1-containing complexes (52–54), the DNA synthesome and replication complexes (55, 56), and the chromatin remodelling LARC, BRM-SIN3A, and ALL-1 supercomplexes (57–59). Overall, these enriched complexes illustrate the general principle that RNA processing, chromatin remodelling, and DNA replication and repair are key effectors of R-loop biology. As further support for the validity of the dataset it is notable that many genes emerging as RLBPs have published roles in R-loop biology, such as subunits of the Mre11-Rad50-Nbs1 complex (15), the RPA complex (60), the BAF complex (61, 62), and the THO complex (63). Thus, our machine learning approach, supported by the CRISPR-FACS screen, recovers known and predicts novel R-loop binding and regulating proteins.

Within the CRISPR screen, we specifically observed that a large and unexpected group of genes functioning in mitochondria increased S9.6 staining, especially mitochondrial ribosomal proteins (**Fig. 4** and **5**). As such, we first set out to test directly if inhibiting mitochondrial translation influenced S9.6 staining. Treatment of cells with the inhibitor tigecycline (64, 65) showed a significant increase in nuclear S9.6 staining (**Fig. 5B**). Indeed, some of our top hits, such as in the mitochondrial degradosome proteins SUPV3L1 and PNPT1, have previously been linked to suppressing R-loop accumulation, both of which are predicted RLBPs and emerge as hits in the CRISPR-FACS screen (66). As such we conclude the mitochondrial enrichment in our CRISPR-FACS experiments do not necessarily represent a technical artifact but instead are associated with *bona fide* increases in S9.6 staining.

**Figure 5.**
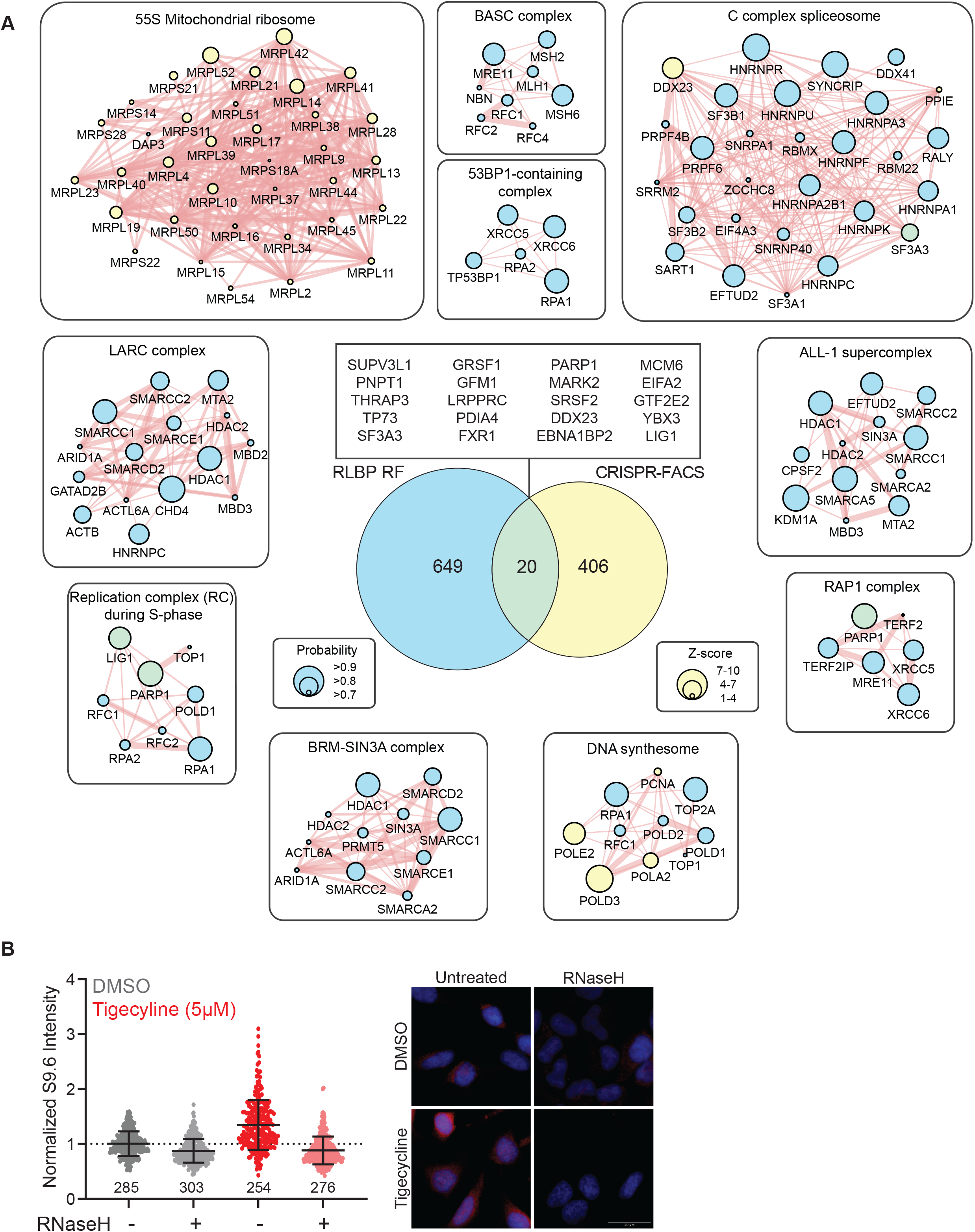
Integrative analysis of enriched complexes involved in R-loop binding and modulation. (A) Network analysis on the combined list of unique random forest predicted RLBPs and genome-wide CRISPR screen enriched genes reveals multiple complexes involved in R-loop biology (p-value < 0.01). Red lines indicate physical interactions between proteins (Cytoscape, Genemania). (B) *Left:* Quantification of immunofluorescence data measuring S9.6 nuclear signal in HeLa cells treated with tigecycline (5uM, 72 hours) cells and DMSO controls (mean +/- SD, One-way ANOVA, N = 3, number of nuclei analyzed is presented under each distribution). *Right:* Representative images.

### Validation of novel candidate R-loop regulating proteins

Intersection of the R-loop modulators identified by our CRISPR screen to our RLBP dataset revealed a small but high-confidence dataset of 20 R-loop modulatory/interacting proteins (**Fig. 5A, middle**). Among the genes that overlap between the RLBP and CRISPR-FACS datasets are known R-loop regulators involved in RNA processing, such as SF3A3, and SRSF2 (67–69), DNA replication and repair proteins like MCM6 and PARP1 (70, 71), and mitochondrial RNA regulators like PNTP1 and GRSF1(66). Given the many literature connections to R-loops within this small overlapping group, we consider it a high confidence set of genes. To validate this classification we chose two additional genes, FXR1 and LIG1, for further study. LIG1 is the major replicative DNA ligase, and a mutant of its yeast orthologue CDC9 was synthetic lethality with RNaseH mutants, and raised S9.6 staining in our previous work (15). FXR1 on the other hand is an RNA binding protein and orthologue of the fragile X and mental retardation gene FMR1(72).

To confirm that these candidate proteins could interact with R-loops, we performed a proximity ligation assay in HeLa cells to quantify the co-localization of FXR1 or LIG1 at sites of S9.6 staining (**FIg. 6A, 6B and S5B)**. In accordance with our data, we were able to observe FXR1-S9.6 and LIG1-S9.6 co-localization in the form of distinct nuclear foci under native conditions. In both cases, the nuclear signal was reduced when the cells were treated with RNaseH, RNaseIII and RNaseT1.

**Figure 6.**
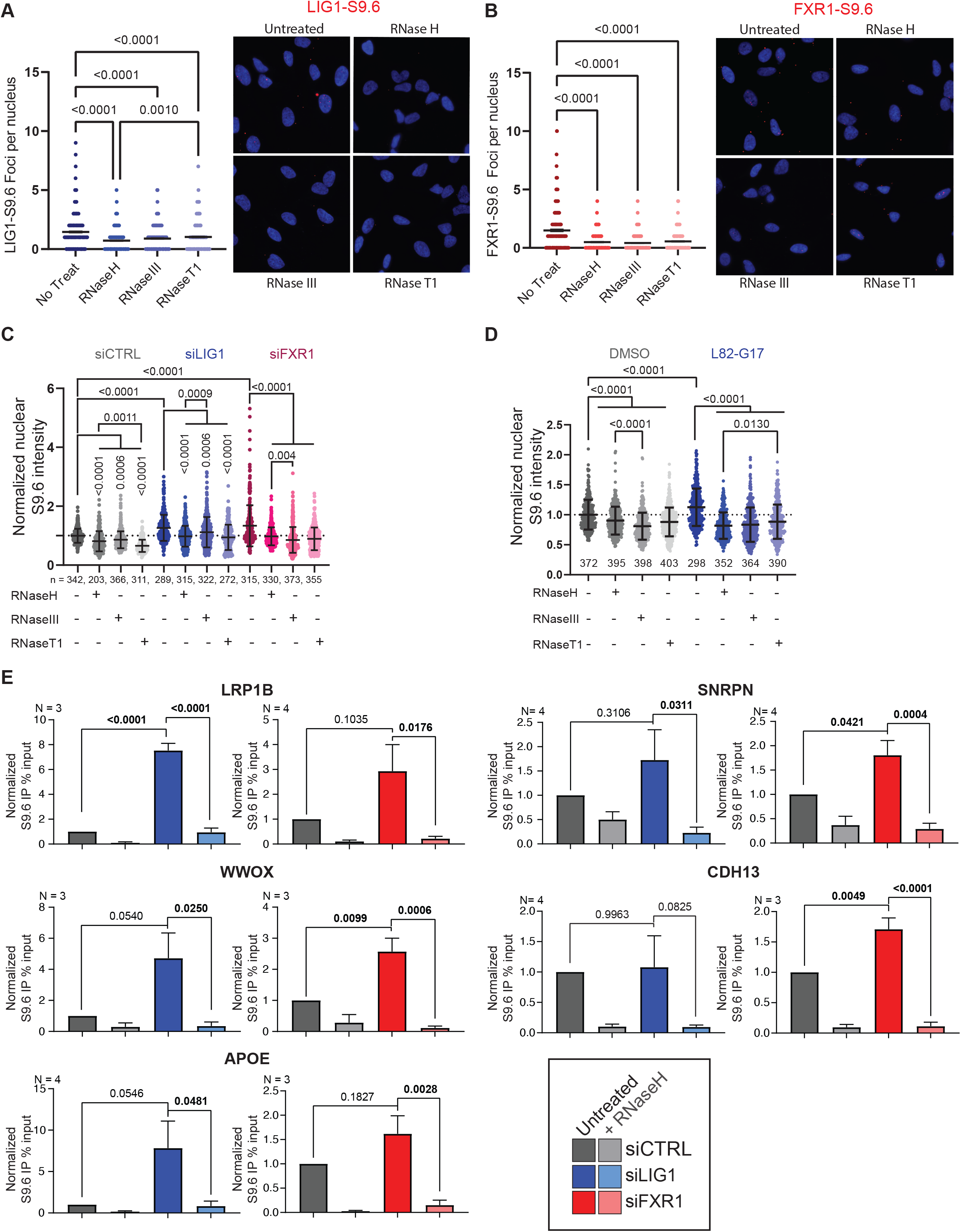
Validation of novel candidate R-loop moduatory proteins. (A-B) *Left:* Quantification of nuclear proximity ligation assay foci showing localization of LIG1 (A) or FXR1 at sites of S9.6 staining. *Right:* Representative images. (One-way ANOVA, N = 3, >300 nuclei analyzed). Scale bars = 20µm). (C) Quantification of nuclear S9.6 immunofluorescence signal in siRNA knockdown of LIG1 (blue) and FXR1 (red). Values are normalized to siCTRL. (mean +/- SD, One-way ANOVA, N = 3, number of nuclei analyzed is presented under each distribution). Representative images can be found in Figure S5C. (D) Quantification of nuclear S9.6 signal from immunofluorescence showing inhibition of LIG1 with L82-G17 (25µM, 4 hours) results in increased nuclear S9.6 staining (mean +/- SD, One-way ANOVA, N = 3, number of nuclei analyzed is presented under each distribution). Representative images can be found in Figure S5D. (E) Relative DRIP-qPCR signal values at *LRP1B, SNRPN, WWOX, APOE* and *CDH13* genes in HeLa cells transfected with indicated siRNAs and treated with *in vitro* RnaseH pre-immunoprecipitation where indicated (mean +/- SEM, One-way ANOVA, at least three independent replicates). Statistically significant values shown in bold.

siRNA depletion of FXR1 or LIG1 in Hela cells led to an increase in nuclear S9.6 staining in both knockdowns compared to the control (**Fig. 6C**). Treatments with RNaseH, RNaseIII and RNaseT1 partially rescued the signal in all datasets, confirming the unspecific nature of the S9.6 antibody. The increase in S9.6 signal in the siLIG1 sample was mostly rescued by RNaseH and RNaseT1, suggesting that knockdown of LIG1 may lead to increased R-loops as well as the accumulation of ssRNA structures within the nucleus. On the other hand, knockdown of FXR1 was partially rescued by RNaseH and significantly reduced by RNaseIII and T1, suggesting that loss of FXR1 may lead to increased levels of R-loops and perhaps to a larger extent dsRNAs and ssRNAs. Additionally, we sought to selectively disrupt LIG1 function using the inhibitor L82-G17, which is an uncompetitive inhibitor of the third step of the ligation reaction phosphodiester bond formation (73). Inhibition of LIG1 by L82-G17 resulted in an increased nuclear S9.6 staining (**Fig. 6D**) that was significantly reduced by RNaseH treatment, suggesting an increase in R-loops. However, RNaseIII and RNaseT1 treatments also reduced the signal intensity, suggesting an increase in the accumulation of dsRNA and ssRNAs species respectively.

To further confirm that LIG1 and FXR1 knockdown cells accumulate more R-loops and to partly circumvent the limitations associated with the non-specific nature of immunofluorescence using the S9.6 antibody, we performed DNA:RNA immunoprecipitation followed by qPCR (DRIP-qPCR) at R-loop prone loci *SNRPN, APOE* and *RPL13A*(16, 19, 74–76), as well as the common fragile sites *WWOX, LRP1B* and *CDH13* (77, 78). *As seen in* ***Figure 6E***, *depletion of LIG1 by siRNA resulted in a significant increase at the LRP1B* locus, and strong increases in signal albeit not statistically significant at the *WWOX* and *APOE* loci. On the other hand, depletion of FXR1 by siRNA led to significant increases in DRIP-qPCR signal at the *WWOX, CDH13* and *SNRPN* loci. No changes were detected at the *RPL13A* locus under either conditions (**Fig. S5E**). All of the changes in DRIP signal observed could be rescued by *in vitro* RNaseH treatment, confirming that the signal was driven by DNA:RNA hybrids. This data suggests that R-loops form at a higher frequency in LIG1 and FXR1 depleted cells, and this occurs at common (e.g. *WWOX*) and distinct genomic sequences (e.g. *APOE, LRP1B, CDH13* and *SNRPN*). Ultimately, we conclude that LIG1 and FXR1 regulate R-loop accumulation and localize to sites of DNA:RNA hybrids, providing independent validation of our high confidence RLBP predictions.

## DISCUSSION

R-loops have emerged in the past decade as pervasive genome-associated structures with roles in gene expression, RNA processing, telomere maintenance, DNA repair and epigenetic regulation. The literature suggests that many dozens of proteins likely regulate R-loop formation, function, and dissolution. Here we reasoned that known R-loop binding proteins will have conserved features that could be used to implicate new proteins in R-loop biology and help assign each protein in the entire human proteome a score that predicts the ability to bind to R-loops. Our machine learning approach robustly predicts RLBPs with functional roles across all of the processes impacted by R-loops. We envision this list as a resource that will interface with the vast and growing number of functional genomic screens taking place in human cell models. For example, screens that capture new genome stability or epigenetic regulators could cross-reference the RLBP dataset to determine if R-loop changes are likely mechanisms of action. We can retrospectively state that this approach would be useful. For example, we have previously shown an R-loop mechanism of genome instability when splicing factors like FIP1L1 or EFTUD2 homologues are mutated, both of these genes appear on our RLBP list (79, 80). More recently, our group and others have found that members of the BAF chromatin remodelling complexes play roles in preventing R-loop associated replication stress (61, 62) and our RLBP list predicts that both catalytic subunits and five accessory BAF subunits are R-loop binding candidates. Additional hits include the DDX23 helicase that plays a crucial role in the regulation of RNAPII pausing to prevent harmful R-loop accumulation (81), PARP1 (also identified in Cristini et al.) which functions in the protection against R-loop associated DNA damage through its interactions with DHX9 and TonEBP(70), and more. In general then, previously recognized R-loop regulators, including those not included in the training set, score very highly in our analysis.

Our machine learning algorithm is built to reduce false positives, and therefore is expected to have an increased false negative rate resulting in potentially missed RLBPs. Features such as the ability to bind nucleic-acids and abundance solely rely on annotated databases that are updated frequently.

Furthermore, large-scale annotations are not entirely accurate. For example, two proteins RnaseH1 and RnaseH2A that bind and resolve R-loops (82) are not classified as “nucleic-acid binding” within the Uniprot database. Each protein scores a probability of 0.103 and 0.346 in our machine learning model, respectively (**Table S2**). However, upon changing their nucleic-acid binding capability to “true” (or 1), their scores change to 0.148 and 0.747 (**Table S2**) respectively where RnaseH2A crosses the threshold to be now called a potential RLBP. The answer to why RnaseH1 is still not called a potential RLBP lies in other features such as abundance. This indicates that with constant updates, better annotations, and more features being discovered, our approach can be reapplied to increase prediction accuracy.

As part of our validation we used the S9.6 DNA:RNA hybrid antibody, an imperfect but useful reagent that has dominated the R-loop field, to profile a large pool of CRISPR knockouts and identify new proteins that increase S9.6 staining. This analysis was clouded by a dramatic enrichment of mitochondrial gene knockouts that increase S9.6 staining. The mitochondrial bias observed in CRISPR screen data is likely multifactorial since our FACS assay measures total S9.6 signal. Due to the cross-reactive nature of the S9.6 antibody, it is likely that S9.6-reactive structures within the mitochondria may be recognized and drive an enrichment of the signal. This is not surprising, as mitochondrial DNA replication can initiate with the formation of an R-loop, and the disruption of mitochondrial R-loop homeostasis likely impacts the readout of our CRISPR screen (66, 83). Accordingly, components of the mitochondrial degradosome complex, PNPT1 and SUPV3L1, were among the most enriched genes associated with an increase in S9.6 staining in our CRISPR and RF screens. These proteins have been directly associated with the suppression of harmful mitochondrial R-loops, supporting the connection between mitochondrial dysfunction and R-loop homeostasis, as well as validating the hits from our screen (84). Furthermore, it is possible that mitochondrial dysfunction may result in altered R-loop dynamics within the cell. In fact, the increase in nuclear S9.6 staining we observed in cells treated with tigecycline, an inhibitor with broad effects on the mitochondria including the induction of ROS-related stress, supports this hypothesis (64, 65). This is in agreement with the idea that ROS induces the formation of RNA:DNA hybrids within the nucleus (85), though the underlying mechanisms of this connection remain unclear. Despite the strong representation of mitochondrial genes, our CRISPR screen also identified genes enriched in pathways relating to R-loop homeostasis like DNA repair. For example, RNaseH2A and ERCC5 which both have well-defined roles in R-loop processing and genome integrity maintenance were identified by our assay (82, 86–92).

LIG1 is an ATP-dependent DNA ligase with functions in DNA replication, recombination and base excision repair, and it was previously shown that LIG1 plays a role in the repair of DNA double strand break lesions associated with R-loop mediated genome instability (93). Here we show that disruption of LIG1 with RNA interference or with chemical inhibition results in an increase in S9.6 staining and DRIP-qPCR, confirming a functional role in R-loop modulation. However, it is not clear why loss of LIG1 would increase R-loops across the genome. LIG1 plays an essential role in completing synthesis of the lagging strand during DNA replication (94), and cells depleted for LIG1 recruit 10 times more PARP in S-phase to unligated Okazaki fragments in order to recruit alternative ligases (95). Furthermore, LIG1 depletion leads to incomplete Okazaki fragment maturation, which in turn affects ATAD5-dependent unloading of PCNA from chromatin (96, 97). Previous work has shown that PCNA accumulation on chromatin due to ATAD5 loss results in transcription-replication conflicts that lead to R-loop formation behind the replication fork. It is possible that interfering with this LIG1-ATAD5-PCNA pathway could result in increased R-loop formation due to retention of excess PCNA on DNA.

FXR1 is an RNA binding protein that interacts with the functionally-similar protein FMR1, which has a proposed role in physically recruiting the R-loop processing helicase DHX9 to prevent harmful R-loop accumulation (98). In addition, both FXR1 and FMR1 are mutated in the R-loop associated disease, Fragile X syndrome (99–101). Here, we show that disruption of FXR1 by RNA interference results in an increase in nuclear S9.6 signal by imaging and DRIP-qPCR which can be partially attributed to R-loops. In addition, we also show that FXR1 co-localizes to sites of S9.6 by PLA, suggesting that FXR1 can be recruited to sites of R-loops. Though the mechanism by which FXR1 impacts R-loop homeostasis remains to be uncovered, we speculate that FXR1 may function in a similar fashion as its orthologue FMR1 as a “detector” of R-loops. Additionally, FXR1 can physically interact with MRE11, which our group has previously identified as important for R-loop suppression by the Fanconi Anemia pathway (15). This FXR1-MRE11 interaction was shown to be particularly important in the cellular response to ROS (102), reinforcing a potential link between mitochondrial dysfunction and R-loop homeostasis. Future work on this previously unrecognized role of FXR1 in R-loop homeostasis may shed light on the mechanisms underlying the genome instability phenotypes associated with Fragile X syndrome and other R-loop associated trinucleotide expansion diseases.

R-loops functioning as mediators of epigenetic states, in telomere regulation, in DNA repair, and at transcription-replication conflicts mean that they have the potential to be recognized and regulated by many proteins. While dozens of proteins have been ascribed functions in R-loop regulation, the complete spectrum of potential R-loop binding and regulating proteins is not understood. This complexity was the ultimate rationale for generation of the RLBP resource presented here. Researchers suspecting a relationship between their protein of interest and R-loop biology can use the dataset presented here as a preliminary investigation tool. Moreover, the emergence of S9.6 alternatives that can detect R-loops more specifically will improve researchers ability to assess R-loop binding and modulatory functions in the future. For example, the recent use of a purified recombinant catalytically inactive human RNase H1 tagged with GFP shows strong specificity against R-loops (103). Refinements to methods and curation of functionally validated RLBPs will help to improve this resource going forward.

## DATA AVAILABILITY

Further information and requests for resources and reagents should be directed to and will be fulfilled by the Lead Contact, Peter Stirling

## ACKNOWLEDGEMENTS

We thank Maxwell Libbrecht, his laboratory, and Dana Bazazeh, for helpful discussions related to the machine learning algorithm. We thank David Huntsman and members of his group for help in preparation of lentiviral stocks. We thank James Wells and members of the Stirling lab for technical assistance.

## SUPPLEMENTARY INFORMATION

**Figure S1**. (A-F), Aliphatic index (A),percentage of positive and negatively charged amino acids (or AAs) (B) and (C), disorder percentage (D), PScore (E), and phosphosite count (F) were compared for the indicated whole proteome, nuclear proteome, and RLBP datasets. The exact p-values resulting from a Mann-Whitney-Wilcoxon test after Bonferroni correction for multiple comparisons are reported.

**Figure S2**. (A-I), (A) Length, (B) hydrophobicity and hydrophilicity (GRAVY), (C) aliphatic index, (D) percentage of low-complexity regions, (E) PScore, (F) disorder percentage, (G) abundance, (H) phosphosite count, and (I) solubility were compared for the indicated Not Predicted (NP) and Uniquely Predicted (UP) datasets of nucleic-acid binding proteins. The exact p-values resulting from a Mann-Whitney-Wilcoxon test after Bonferroni correction for multiple comparisons are reported.

**Figure S3**. (A) Venn diagram comparing proteins shared amongst newly predicted candidate RLBPs, and the independent validation dataset.

**Figure S4**. (A) FACS histogram of HeLa cells stained with the S9.6 antibody. Populations depicted represent unstained cells (orange), native S9.6 staining (blue) and camptothecin-treated (red - 10μM, 2 hours). B) FACS distribution of HeLa stained with the S9.6 Antibody with representative gates used for TKOv3 CRISPR screen FACS sorting. (C) Sample readcount data of the top enriched sgRNA (chr2:55907992-55908011_PNPT1_-) against PNPT1 across the different sorted population populations (log10 scale, N=2). (D) Venn diagram of overlap between our genome-wide CRISPR screen enriched genes (unique) and the Cristini and Wang datasets.

**Figure S5**. (A) Western blot of siRNA knockdowns. (B) Quantification (*Left*) and representative images (*Right*) of PLA antibody controls). (C) Representative images of S9.6 staining from the quantification data presented in Figure 6B (siRNA knockdown). (D) Representative images of S9.6 staining from the quantification data presented in Figure 6C (LIG1 inhibition with L82-G17). (E) Relative DRIP-qPCR signal values at *RPL13A* genes in HeLa cells transfected with indicated siRNAs and treated with *in vitro* RNaseH pre-immunoprecipitation where indicated (mean +/-SEM, One-way ANOVA, N = 5).

**Table S1**. Proteins found to interact with R-loops by LC/MS methods. *Sheet 1*: R-loop binding proteins (RLBPs). *Sheet 2*: PFAM domain enrichment values for RLBPs.

**Table S2**. Protein features for the human proteome. *Sheet 1*: All features used for the random forest algorithm per protein. *Sheet 2*: PFAM domains per protein for the human proteome.

**Table S3**. Probabilities for each protein called by the easy ensemble random forest algorithm. *Sheet 1*: Probability for each protein in all 100 models. *Sheet 2*: Averaged probability for each protein. *Sheet 3*: Performance metrics on the test set. *Sheet 4*: Candidate RLBPs with probabilities greater than a threshold of >=0.7. *Sheet 5*: GO Pathway enrichment values for uniquely predicted candidate RLBPs.

*Sheet 6*: Probability for RnaseH1 and RnaseH2A.

**Table S4**. CRISPR Screen differential analysis by DESEQ2. *Sheet 1:* DESEQ2 scores for all sgRNAs. *Sheet 2:* DESEQ2 scores for enriched sgRNAs in the S9.6 high fraction (hit list).

**Table S5**. Oligonucleotides used in this study.

