## Supplemental Figures S1-S4 for "Global prediction of candidate R-loop binding and R-loop regulatory proteins"

RLBP Supplemental Fig 1.

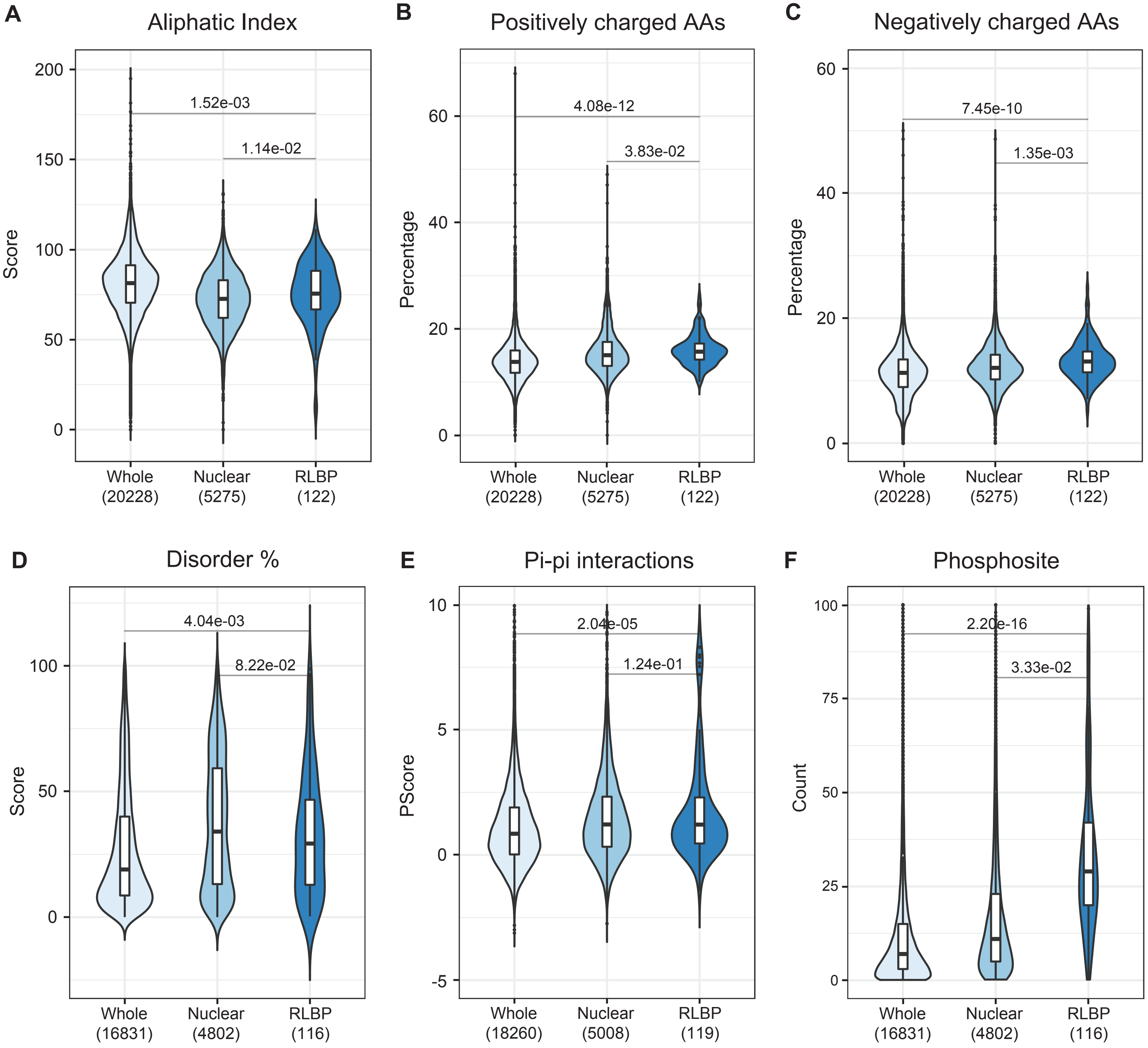

RLBP Supplemental Fig 2.

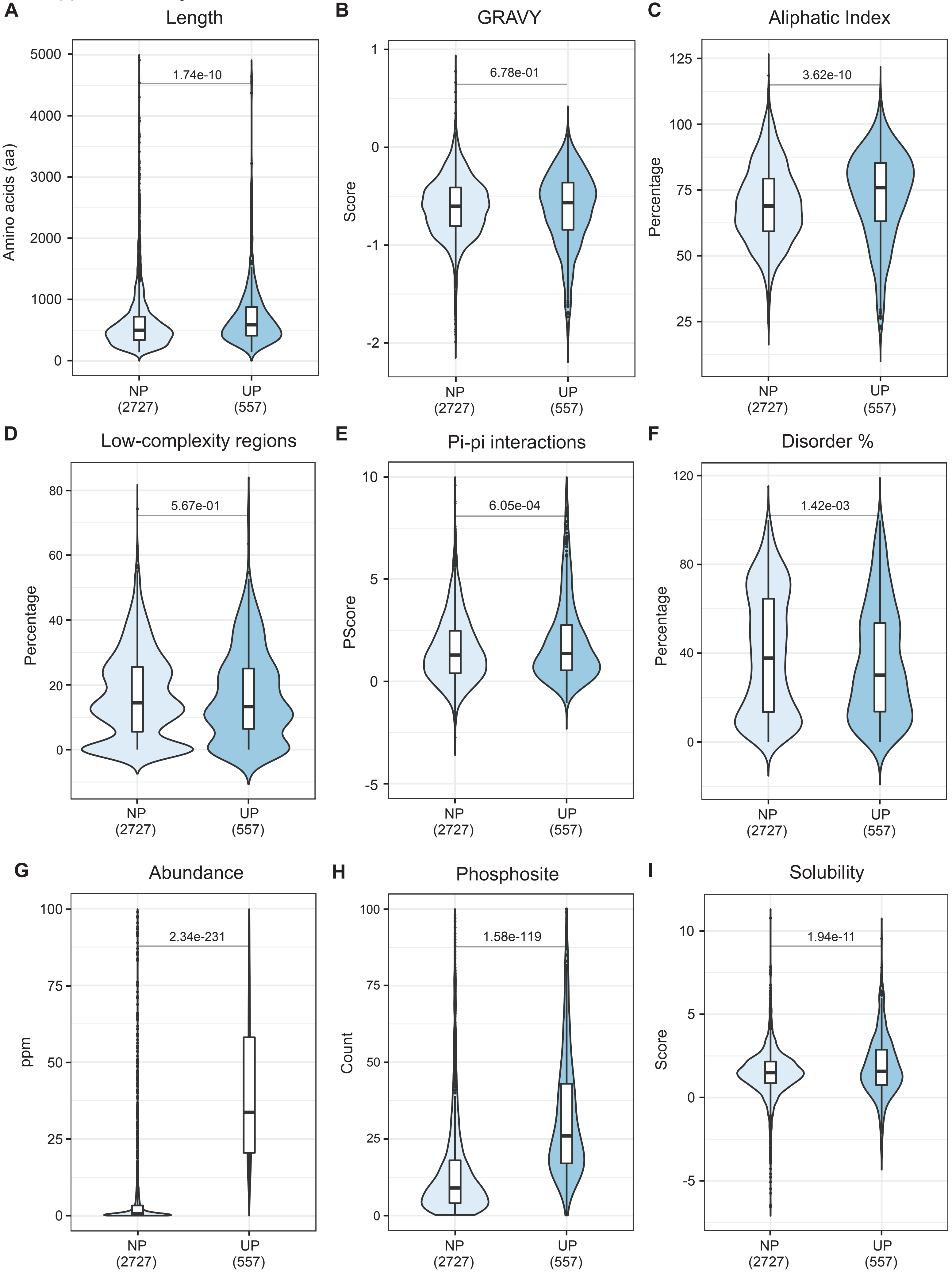

RLBP Supplemental Fig 3.

**A**

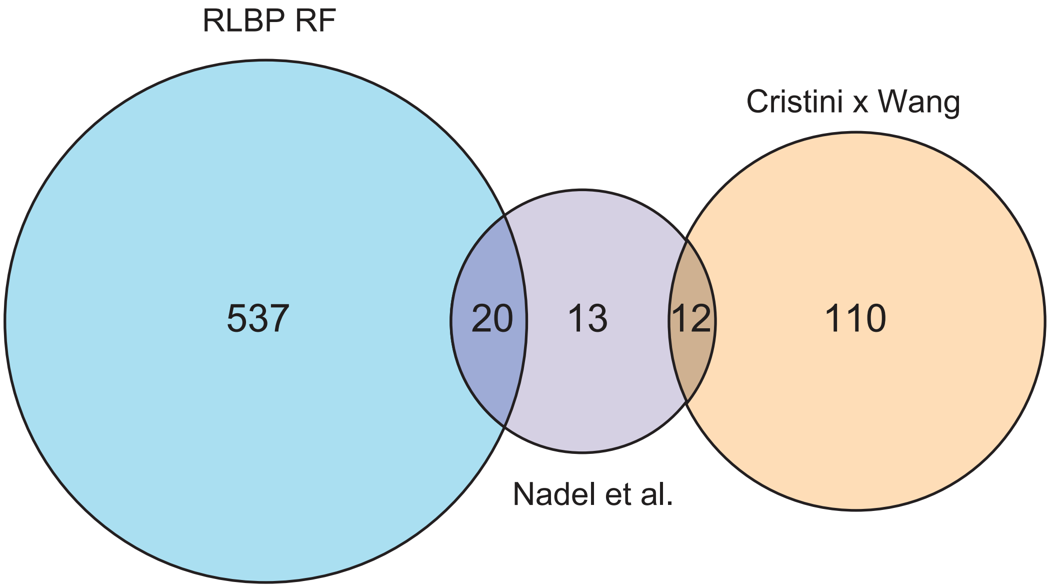

RLBP Supplemental Fig. 4

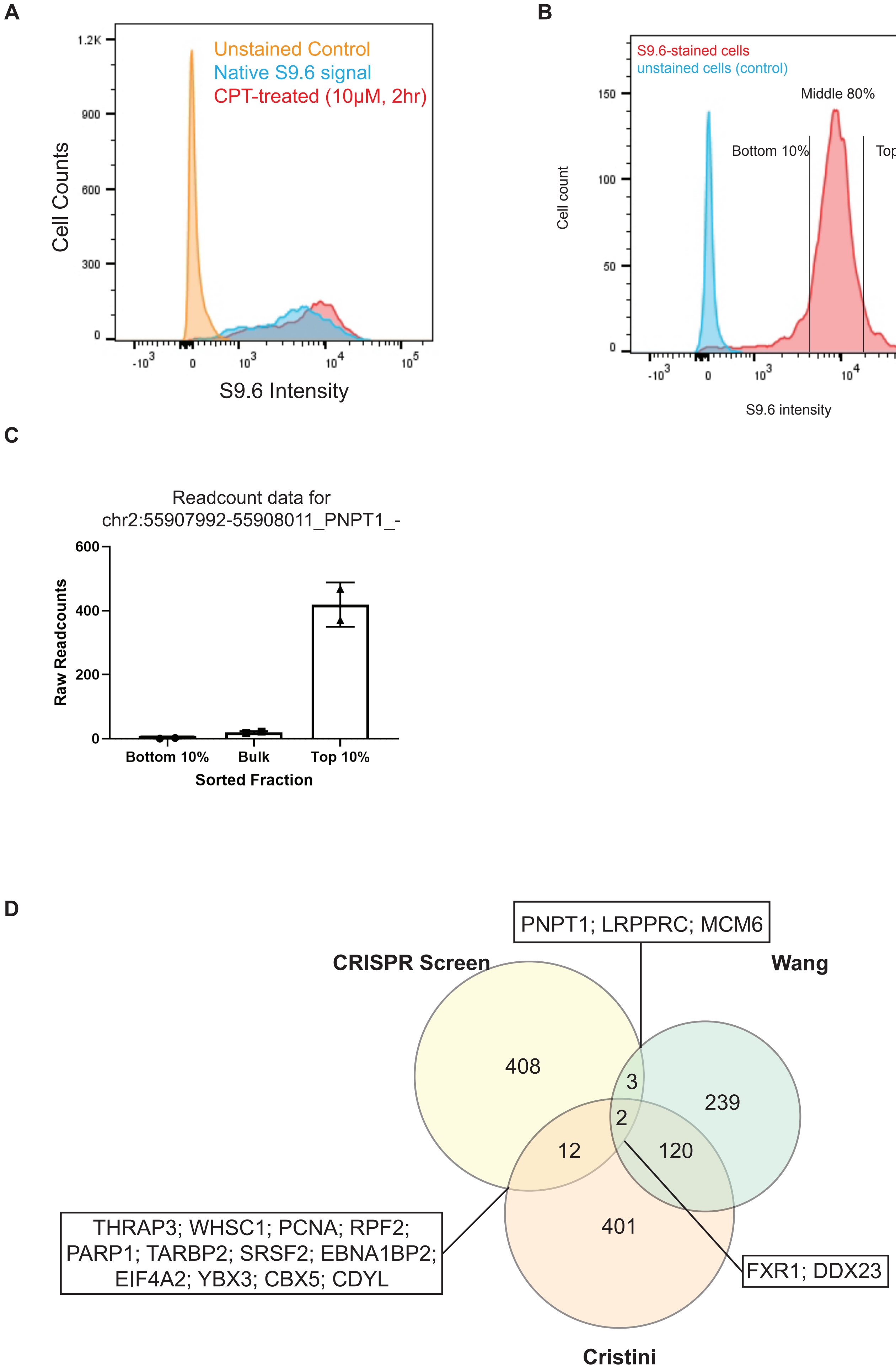

Supplemental Figure 5.

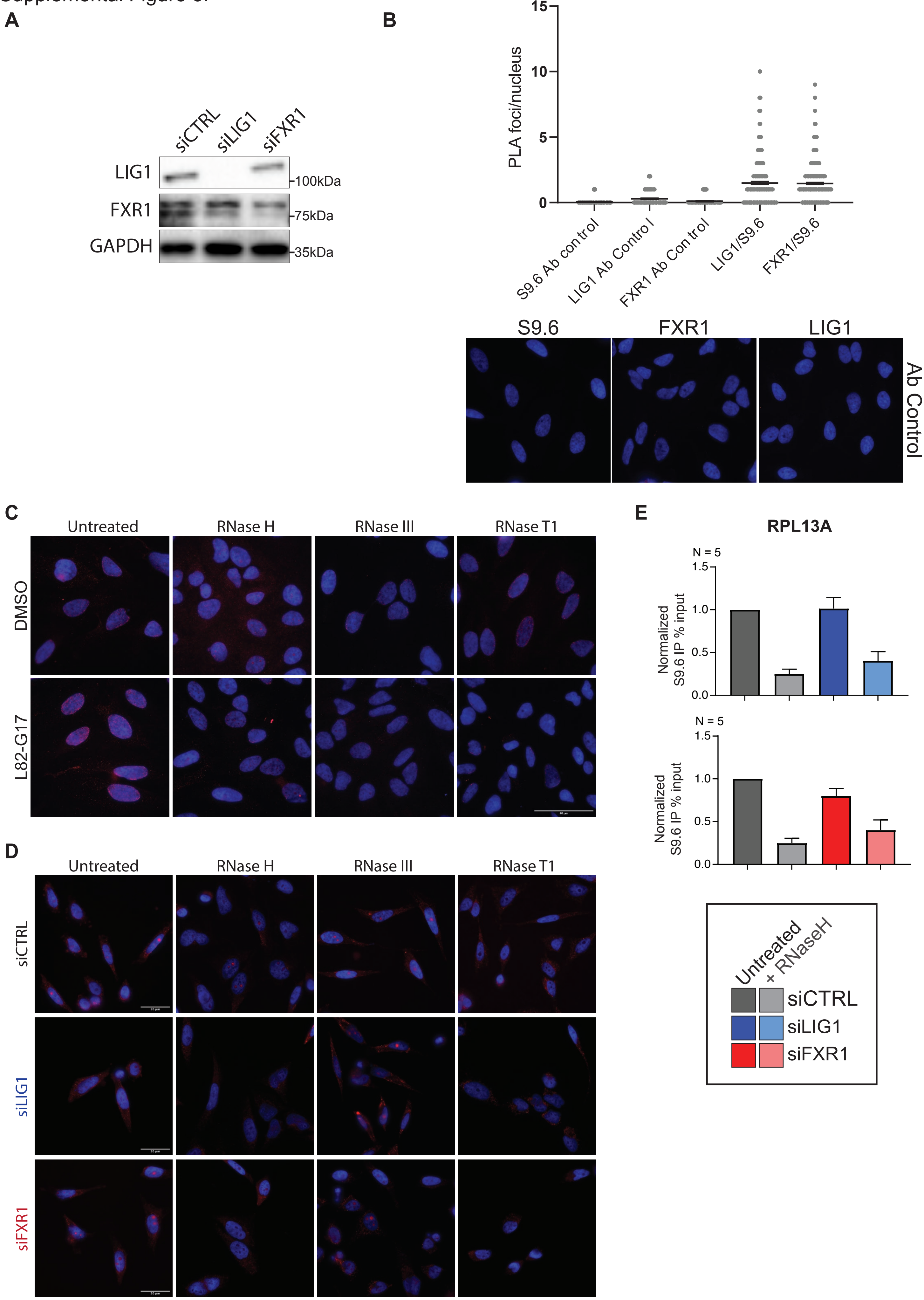
